# Rad52 sorts and stacks Rad51 at the DNA junction to promote homologous recombination

**DOI:** 10.1101/2024.11.07.622519

**Authors:** Jaigeeth Deveryshetty, Ayush Mistry, Sushil Pangeni, Mohamed Ghoneim, Monica Tokmina-Lukaszewska, Vikas Kaushik, Angela Taddei, Taekjip Ha, Brian Bothner, Edwin Antony

**Author notes:** Address correspondence to: Edwin Antony, Department of Biochemistry and Molecular Biology, St. Louis University School of Medicine, St. Louis, MO 63104. Authors contributed equally to this study (Jaigeeth Deveryshetty, Ayush Mistry).

## Abstract

Homologous recombination (HR) repairs double-stranded DNA breaks (DSBs). The DSBs are resected to yield single-stranded DNA (ssDNA) that are coated by Replication Protein A (RPA). Rad51 is a recombinase and catalyzes strand invasion and the search for homology. However, it binds to ssDNA with lower affinity than RPA. Thus, mediator proteins such as Rad52/BRCA2 are required to promote Rad51 binding to RPA-coated ssDNA, but the underlying mechanisms remain poorly understood. *Saccharomyces cerevisiae* Rad52 interacts with Rad51 through two distinct binding modes. We here uncover that the Rad51-binding site in the disordered C-terminus of Rad52 (mode-1) sorts polydisperse Rad51 into discrete monomers. The second Rad51 binding site resides in the ordered N-terminal ring of Rad52 (mode-2), but this interaction occurs at only one position on the ring. In single molecule confocal fluorescence microscopy combined with optical tweezer analysis, we directly visualize filament formation using fluorescent-Rad51. Rad52 catalyzes Rad51 loading onto RPA-coated ssDNA, with a distinct preference for junctions, but no filament growth is observed. Deletion of the C-terminus of Rad52 results in loss of Rad51 sorting and abrogates Rad51 binding to RPA-coated DNA. While BRCA2 and Rad52 are structurally unrelated, many of these functional features are conserved. We describe a concerted *Sort & Stack* mechanism for mediator proteins in promoting HR.

## Introduction

Accumulation of DNA damage results in genomic instability and is a hallmark of most cancers^1^. Double-stranded DNA breaks (DSBs) are particularly harmful as they give rise to chromosomal aberrations and deletions^2^. Homologous recombination (HR) is a key DNA repair pathway that protects cells from the harmful genomic instability effects introduced by DSBs^3-5^. In HR, the DNA break in one allele is repaired using the homologous sequence in the undamaged sister chromatid as a template. HR is orchestrated through a sequence of DNA break recognition, processing, homology search & recognition, and resolution steps that are catalyzed by over three dozen proteins^3-5^. In *Saccharomyces cerevisiae*, the DSB is first recognized and resected to yield 3’
s ssDNA overhangs which are subsequently sequestered by the ssDNA binding protein Replication Protein A (RPA) to serve as a nucleoprotein platform for the recruitment of downstream HR enzymes^6,7^. Rad51 is the recombinase that catalyzes homology search and pairs the correct complementary sequences – a process called strand exchange or synapsis. Rad51 binds to the RPA-coated ssDNA (nucleation) and forms an ATP-dependent cooperative helical filament (growth). This pre-synaptic Rad51 filament formation step is a key regulatory event in HR.

The impediment to spontaneous Rad51 binding to the resected ssDNA arises from the high affinity of RPA-ssDNA interactions (K_D_<1 nM) compared to Rad51 binding to ssDNA (K_D_∼1 µM)^8,9^. Thus, several mediator proteins and Rad51-paralogs function to regulate pre-synaptic Rad51 DNA binding/nucleation and filament growth during HR. Pro-HR mediators promote the nucleation and growth of the Rad51 filament on RPA-coated ssDNA whereas anti-HR mediators suppress HR by removing Rad51 molecules from the DNA^3^. In yeast, Rad52 is the key pro-HR mediator and a functional homolog of the human BRCA2 protein^10^. Anti-HR mediators are primarily translocases or helicases that utilize ATP-dependent physical movement on ssDNA to remove Rad51^5,11^. A key anti-HR mediator in yeast is the Srs2 helicase that physically moves on the ssDNA to strip Rad51^11-15^. Rad51 filaments are further stabilized by Rad51 paralogs such as Rad55-Rad57 and the SHU complex made of Csm2-Psy3-Shu1-Shu2 subunits^16^. The paralogs are thought to intercalate or cap the ends of the Rad51 filament and protect them from disassembly by anti-HR mediators such as Srs2^17-20^. While vast efforts have been undertaken to decipher the functions of mediator proteins and Rad51-paralogs in HR, knowledge of where they bind, how they interact with each other and Rad51, and where they are positioned on the DNA are poorly resolved.

In this study, we focus on the mechanism of Rad52-promoted Rad51 filament formation on RPA-coated DNA. We recently uncovered that yeast Rad52 is a homodecameric ring with each subunit possessing an ordered N-terminal and disordered C-terminal half^21^. The N-terminal region promotes oligomerization of Rad52 and ssDNA interactions whereas the C-terminal region harbors the Rad51 and RPA interaction motifs. Interestingly, we also uncovered additional interactions with Rad51 through the ordered N-terminal ring, but these were asymmetric and captured alongside only one Rad52 subunit^21^. Thus, we defined two modes of Rad52-Rad51 interactions: *Mode-1* interactions occur between Rad51 and the disordered C-terminus of Rad52 and *Mode-2* interactions are enacted between Rad51 and the ordered N-terminus of Rad52. But the functional significance of these modes of interactions were not clear.

Several groups have attempted to visualize and capture the dynamics of yeast Rad51 filament formation. Single-molecule DNA curtain assays have been the most useful in deriving information about binding and dissociation of Rad51, Rad52, RPA, and Rad51-paralog proteins during pre-synaptic filament formation on long ssDNA substrates^18,22-24^. However, Rad51 was never directly visualized as attempts to fluorescently label the protein rendered it inactive. Nevertheless, the binding and dissociation signal from fluorescent RPA served as a reasonable proxy for Rad51 filament dynamics^24-28^. Liu *et. al*. recently described a strategy to generate superfolder green fluorescent proteins (sfGFP)-tagged Rad51 (Rad51^GFP^) using flexible linkers engineered within the N-terminal non-conserved region^29^. This protein engineering strategy generated a functionally active version of Rad51 *in vivo*.

Here, using recombinant Rad51^GFP^ and derivative versions, we directly visualize Rad51 binding events in single-molecule high-resolution optical trap-based tweezers and fluorescence imaging. Using structural mass spectrometry and biophysical approaches we define the functions of both modes of Rad52-Rad51 interactions. Mode-1 sorts Rad51 into monomers whereas mode-2 stacks Rad51 at a single position on Rad52. On RPA-coated ssDNA carrying junctions, the Rad52-Rad51 complex preferentially binds close to the ss-dsDNA junction, but no extension or growth of the Rad51 nucleoprotein is observed. Deletion of the C-terminus of Rad52 results in a complete loss of Rad51 binding to the RPA-coated DNA. The work uncovers how Rad52 functions as a mediator protein in pre-synapsis.

## Results

### S. cerevisiae Rad51 is polydisperse in solution

Most recombinases including bacterial RecA and human RAD51 are oligomers in solution and cooperatively interact with DNA to form nucleoprotein filaments^30-32^. Structural studies have revealed the interactions between *S. cerevisiae* Rad51 protomers within a filament in the absence or presence of DNA (**Figures 1A & B**)^33,34^. To obtain a quantitative measure of the polydispersity of yeast Rad51 under our reaction conditions, we assessed the distributions at both low and high concentrations using mass photometry (MP) and analytical ultracentrifugation sedimentation velocity (AUC^SV^), respectively. Both experiments reveal a wide distribution of oligomeric species ranging from monomers to decamers at low (100 nM) or high (10 µM) concentrations (**Figures 1C & D)**. To further understand the nature of contacts between Rad51 molecules, we performed crosslinking mass spectrometry (XL-MS) analysis using suberic acid bis 3-sulfo-N-hydroxysuccinimide ester (BS3) which crosslinks primary amines within a ∼12-24 Å distance^35^. More than 50 crosslinks (XLs) are observed within Rad51 (**Figure 1E**) which represents both intra- and inter-Rad51 contacts. No XLs are identified in the disordered N-terminal region of Rad51 as there are no Lys residues in this region. Moreover, this region is not conserved among eukaryotic Rad51 proteins.

**Figure 1.**
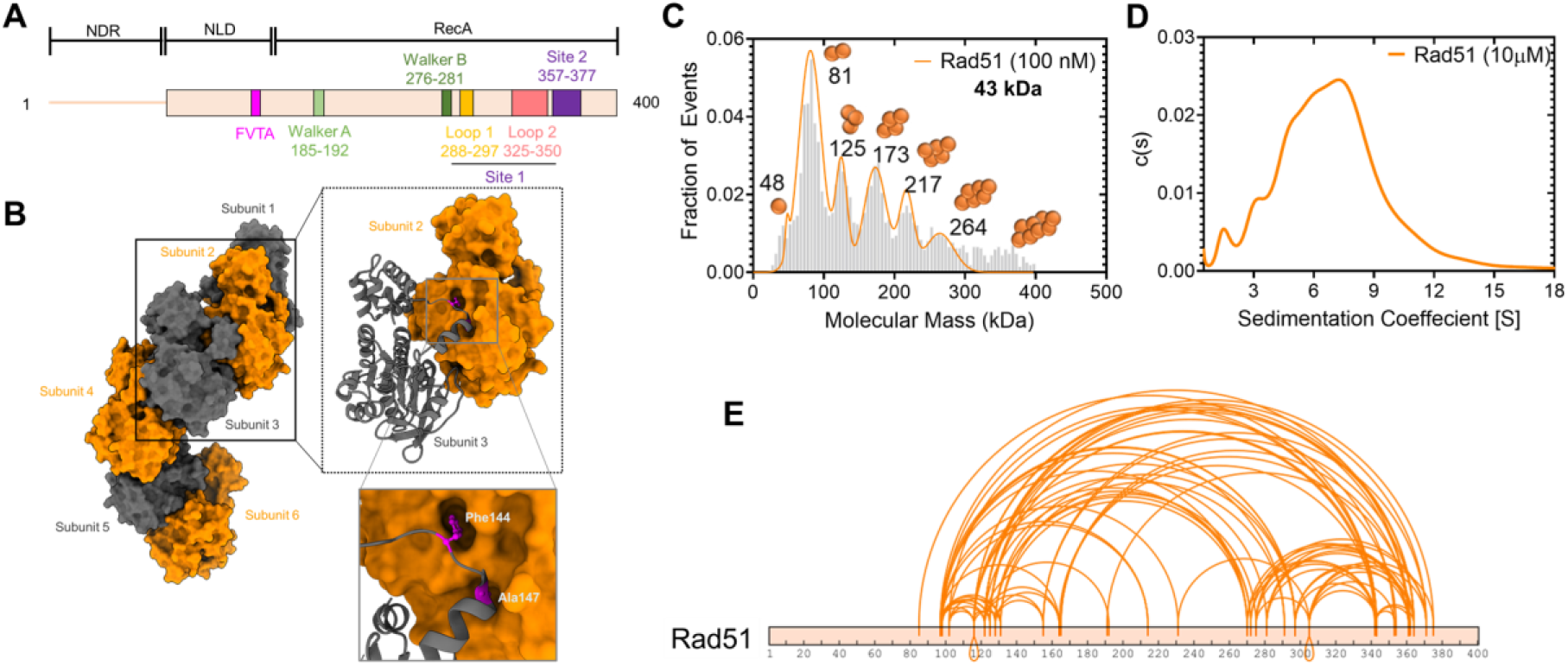
*S. cerevisiae* Rad51 is polydisperse in solution. **A)** Schematic of Rad51 denoting the positions of the Walker A and B motifs for ATP binding and hydrolysis. The FVTA motif promotes Rad51 oligomerization. Loops 1 and 2 in site 1 and residues in site 2 are critical for DNA binding. **B)** Structure of the Rad51 filament in the absence of DNA (PDB:1SZP). The monomeric units are colored grey and orange. Phe-144 and Ala-147 dock into defined pockets of the adjacent Rad51 protomer to promote oligomerization. **C)** Mass photometry analysis of Rad51 (100 nM) shows a wide range of oligomers ranging from monomers through decamers. **D)** Analytical ultracentrifugation sedimentation velocity analysis of Rad51 at higher concentrations (10 µM) shows a similar distribution of oligomers. **E)** Crosslinking mass spectrometry analysis of Rad51 using BS3 crosslinker show extensive intra- and inter-Rad51 crosslinks.

### Crosslinking mass spectrometry reveals interactions between Rad51 and two distinct binding sites in Rad52

Rad52 physically interacts with Rad51 through two motifs in the disordered C-terminus of Rad52 (**Figure 2A**). Using cryoEM, we recently identified another site of interaction between Rad51 and the N-terminal ordered ring of Rad52^21^. Given the lower resolution of the structural data, the precise motifs that promote this interaction were not identified. Thus, to better understand the details of these two modes of binding, we performed XL-MS analysis of the Rad52-Rad51 complex in the absence of DNA. Rad52 alone shows extensive XLs within the N- and C-terminal halves, and between the two halves (**Figure 2B**). These data agree with the structural and hydrogen-deuterium exchange mass spectrometry analysis performed earlier^21^. Since Rad52 is a homodecamer, the complex was formed using a 1:10 molar ratio of Rad52 (decamer):Rad51 so as to saturate all the binding sites. Distinct sets of XLs are captured between Rad51 and the two binding regions in Rad52 (**Figure 2C**). For clarity, we define these as two modes of interactions: *mode-1* and *mode-2*. The mode-1 interaction site(s) resides in the C-terminus of Rad52 and crosslinks to two regions in Rad51 (residues 100-160 and 300-380). In contrast, the mode-2 interaction site in the N-terminal half of Rad52 predominantly interacts with one region in Rad51 (residues 270-380), although two XLs to a region around residue 120 are also observed (**Figure 2C**). It should be noted that BS3 XL-MS only reports on regions that carry primary amines and should be used as a coarse-grain view of the interaction between Rad52 and Rad51. To get a structural perspective of these interactions, we used AlphFold2 (AF2) prediction to generate a model of the Rad52-Rad51 complex (**Figure 2D**). In agreement with our results, AF2 models two regions of contact between Rad52 and Rad51. Mode-1 interactions map to the C-terminal half of Rad52 whereas mode-2 interactions reside in the N-terminal half of Rad52 (**Figure 2D**).

**Figure 2.**
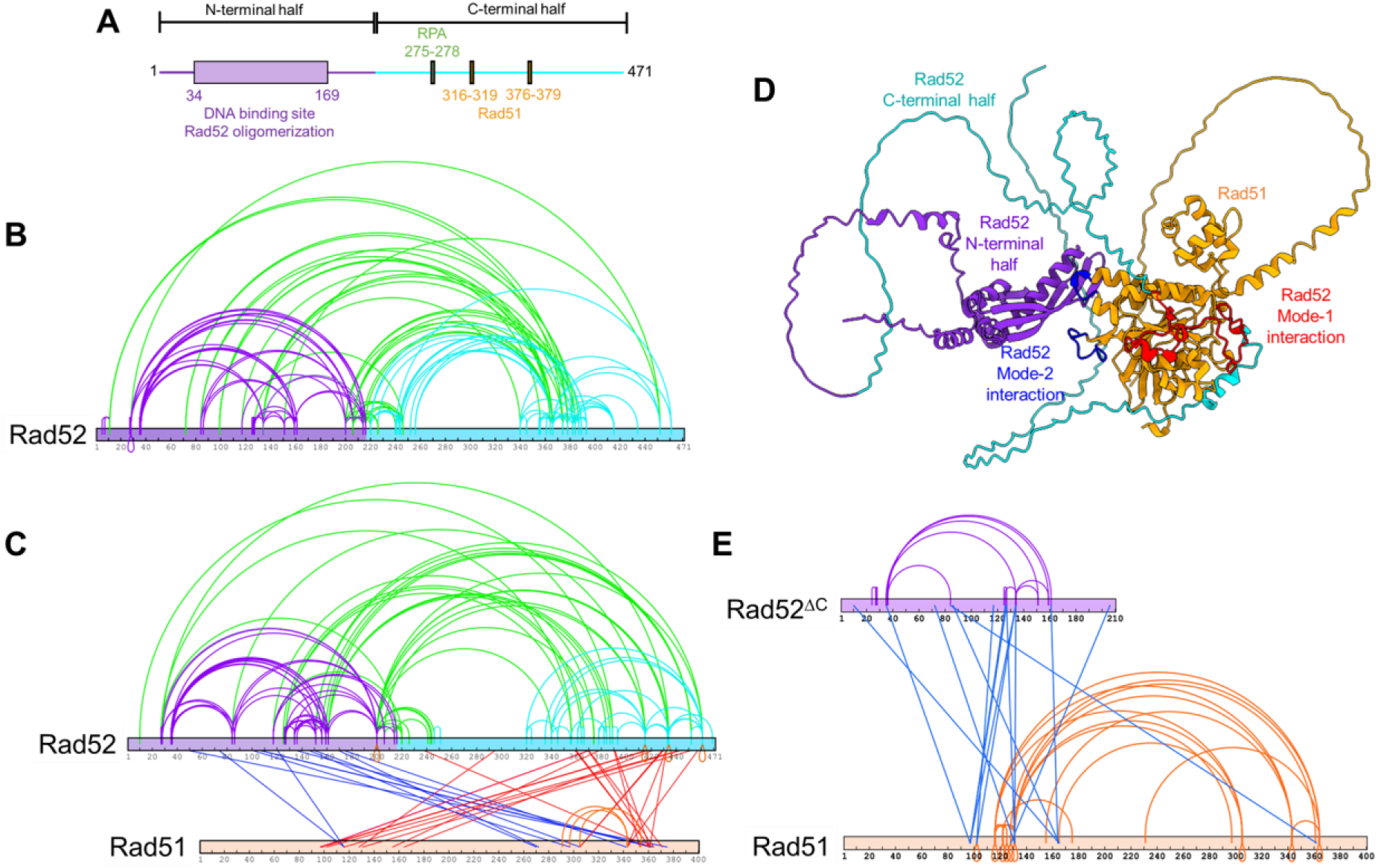
The C-terminus of Rad52 sorts Rad51 into monomers. **A)** Schematic of Rad52 showing the ordered N-terminal and disordered C-terminal halves of Rad52. The RPA and Rad51 binding motifs reside in the C-terminal half. **B)** Crosslinking mass-spectrometry (XL-MS) analysis of Rad52 reveals extensive crosslinks within the N-terminal (purple) and C-terminal (cyan) halves, and between the two halves (green). **C)** XL-MS analysis of the Rad52-Rad51 complex reveals intra-Rad52 crosslinks as shown in panel B, in addition to two sets of crosslinks between Rad51 and Rad52. Crosslinks between the N-terminal half of Rad52 and Rad51 are shown in blue, while crosslinks between the C-terminal half of Rad52 and Rad51 are shown in red. Interestingly, almost all of the inter- and intra-Rad51 crosslinks observed in the Rad51 alone XL-MS analysis are absent when in complex with Rad52. Only three crosslinks within Rad51 are captured (orange). **D)** AlphaFold-2 prediction of the complex between one Rad52 subunit and Rad51. Residues in Rad51 that are proposed to mediate mode-1 interactions with Rad52 are shown in red. Residues that mediate mode-2 interactions are shown in blue and reside in the L1 and L2 loops of Rad51. **E)**. Rad52^ΔC^, lacking the C-terminal half of Rad52 retains interactions with Rad51. XL-MS analysis of this complex reveals the reappearance of the inter- and intra-Rad51 crosslinks suggesting that Rad51 forms oligomers when bound to Rad52^ΔC^. Thus, the Rad51 sorting properties arise from interactions with the C-terminus of Rad52 (mode-1).

### The C-terminus of Rad52 sorts Rad51 into monomers

One striking observation in the XL-MS analysis of the Rad52-Rad51 complex is the loss of almost all intra- and inter-Rad51 crosslinks (**Figure 2C**) which were observed in the XL-MS of Rad51 alone (**Figure 1E**). Only three XLs within Rad51 were captured in the complex (shown in orange in **Figure 2C**) and mapped to the subunit-subunit interface in the Rad51 filament (**Supplementary Figure 1**). In the AF2-model, the Rad51 binding site in the C-terminal half of Rad52 binds across the Rad51-oligomerization interface (**Figure 2D**). This interaction is reminiscent of the interactions between the BRC repeats of human BRCA2 and human RAD51 and promotes monomerization (**Supplementary Figure 2**)^36^. In yeast Rad52, binding mode-1 in the C-terminus is mediated by two known Rad51 interaction patches – 316-FVTA-319 and 337-FDPK-340^37-39^ with Phe-319 and Phe-337 docking into hydrophobic pockets in Rad51 (**Supplementary Figure 2A**). Similar Phe residues are conserved in the BRC4 repeat of BRCA2, but only one was captured in the crystal structure^36^ (**Supplementary Figure 2B**). Thus, we tested whether this region in Rad52 promotes monomerization of Rad51 by deleting the C-terminal half (Rad52^ΔC^). In XL-MS analysis of the Rad52^ΔC^-Rad51 complex crosslinks between the two proteins are observed (**Figure 2E**). Interestingly, the intra- and inter-Rad51 crosslinks reappear in the Rad52^ΔC^-Rad51 complex (**Figure 2E**) suggesting that the Rad51 monomerization (sorting) properties are lost in this mutant. In AUC^SV^ analysis Rad52^ΔC^ binds to Rad51 in agreement with contacts mediated by mode-2 interactions within the N-terminal half of Rad52^21^ (**Supplementary Figure 3**). However, unlike the Rad52-Rad51 complex that produces defined species, the Rad52^ΔC^-Rad51 complex is polydisperse. These data are in agreement with the reappearance of the XLs between Rad51 molecules in the Rad52^ΔC^-Rad51 complex.

To directly assess the Rad51 sorting function of the C-terminus of Rad52, we generated a C-terminal fragment of Rad52 that encompass the two Rad51-binding motifs (Rad52^ΔN*^; **Figure 3A**). Based on the XL-MS analysis, we hypothesized that this fragment would shift the equilibrium of Rad51 in solution from oligomers to monomers (**Figure 3B**). In mass photometry, Rad52^ΔN*^ interacts with Rad51 in a concentration-dependent manner and sorts Rad51 into monomeric units (**Figures 3C-E**, & **Supplementary Figure 4**). These data show that mode-1 interactions are driven by the C-terminal half of Rad52 and function to sort Rad51 into monomers. We had previously shown that Rad51 interacts with the ordered N-terminal ring of Rad52, and this mode-2 interaction occurs in an asymmetric manner within a single subunit in the ring^21^. Thus, we propose a *sort and stack model* where mode-1 interactions sort Rad51 and mode-2 interactions likely deposit the sorted Rad51 molecules at a defined position on Rad52 towards loading on to the resected ssDNA during the pre-synaptic phase of homologous recombination.

**Figure 3.**
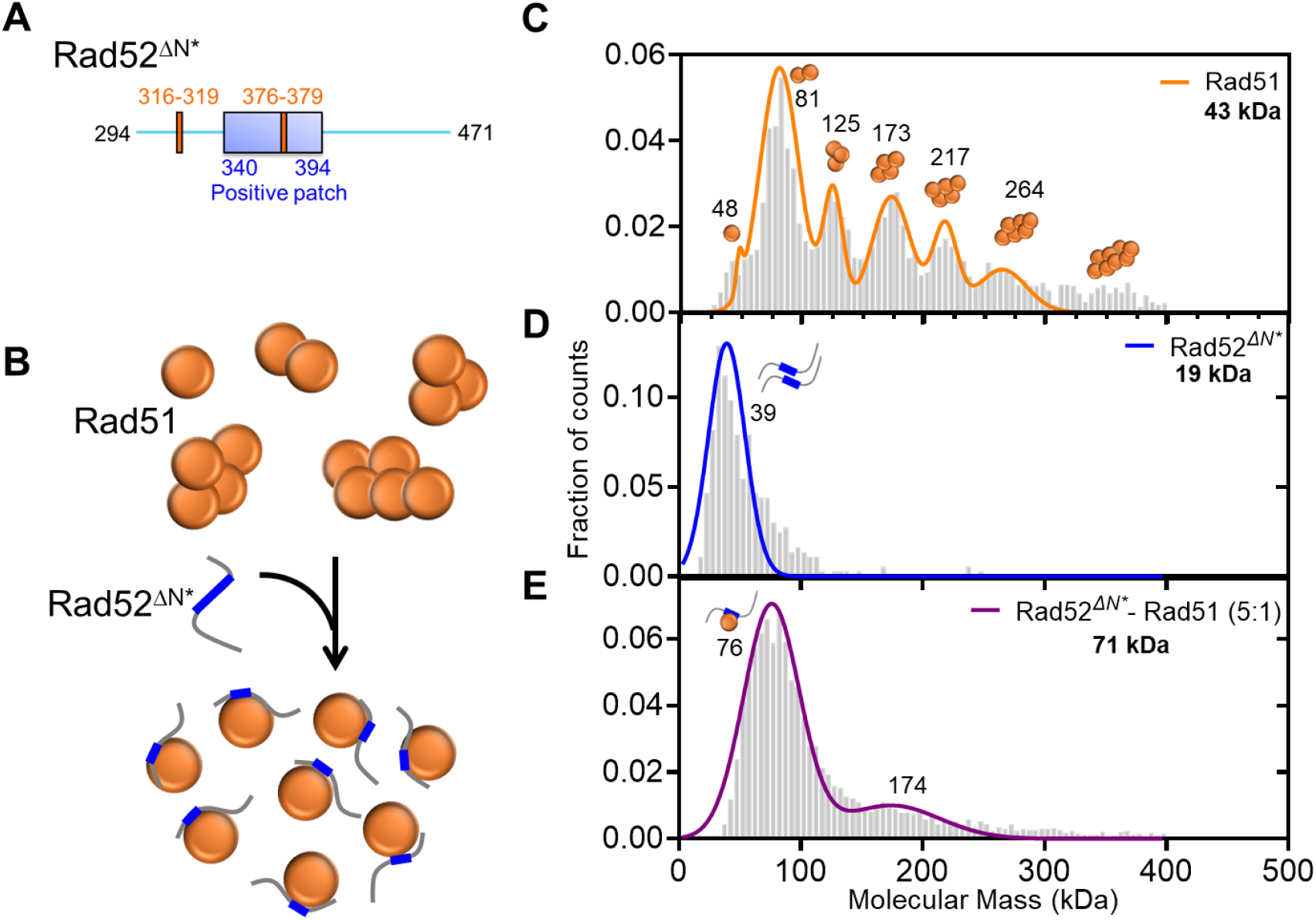
The Rad51-sorting function is mediated through mode-1 interactions in the disordered C-terminus of Rad52. **A)** Schematic of the two Rad51 interaction motifs in the disordered C-terminus of Rad52. The Rad52^ΔN^* is a construct that possesses both these motifs and **B)** is able to sort Rad51 from a polydisperse to monodispersed species in solution. Mass photometry analysis of **C)** Rad51, **D)** Rad52^ΔN*^, and **E)** the Rad51-Rad52^ΔN*^ complex. Rad51 is polydisperse with monomeric to decameric species observed in solution. Rad52^ΔN*^is single lower molecular weight species in solution. It should be noted that the predicted mass is below the detectable limit of the methodology. The Rad51-Rad52^ΔN*^ complex shows a loss of the higher order Rad51 species and accumulation of a predominantly single species that corresponds to one Rad51 molecule bound per Rad52^ΔN*^.

### Direct visualization of Rad51 binding and filament formation on ssDNA

A method to directly visualize and quantitate Rad51 binding onto DNA is required to decipher how mode-1 and mode-2 interactions drive Rad52-promoted Rad51 filament formation. Our many attempts to generate fluorescently labeled yeast Rad51 using traditional approaches (amino-terminal labeling, maleimide-coupling, or through incorporation of non-canonical amino acids) resulted in loss of activity. Recent work by Liu *et. al*. demonstrated that engineering a superfolder green fluorescent protein (sfGFP) at position 55 in the disordered and non-conserved N-terminal region of Rad51 through two additional 16 aa linkers (**Figures 4A & B**) produced a functional recombinase *in vivo*^*29*^.

**Figure 4.**
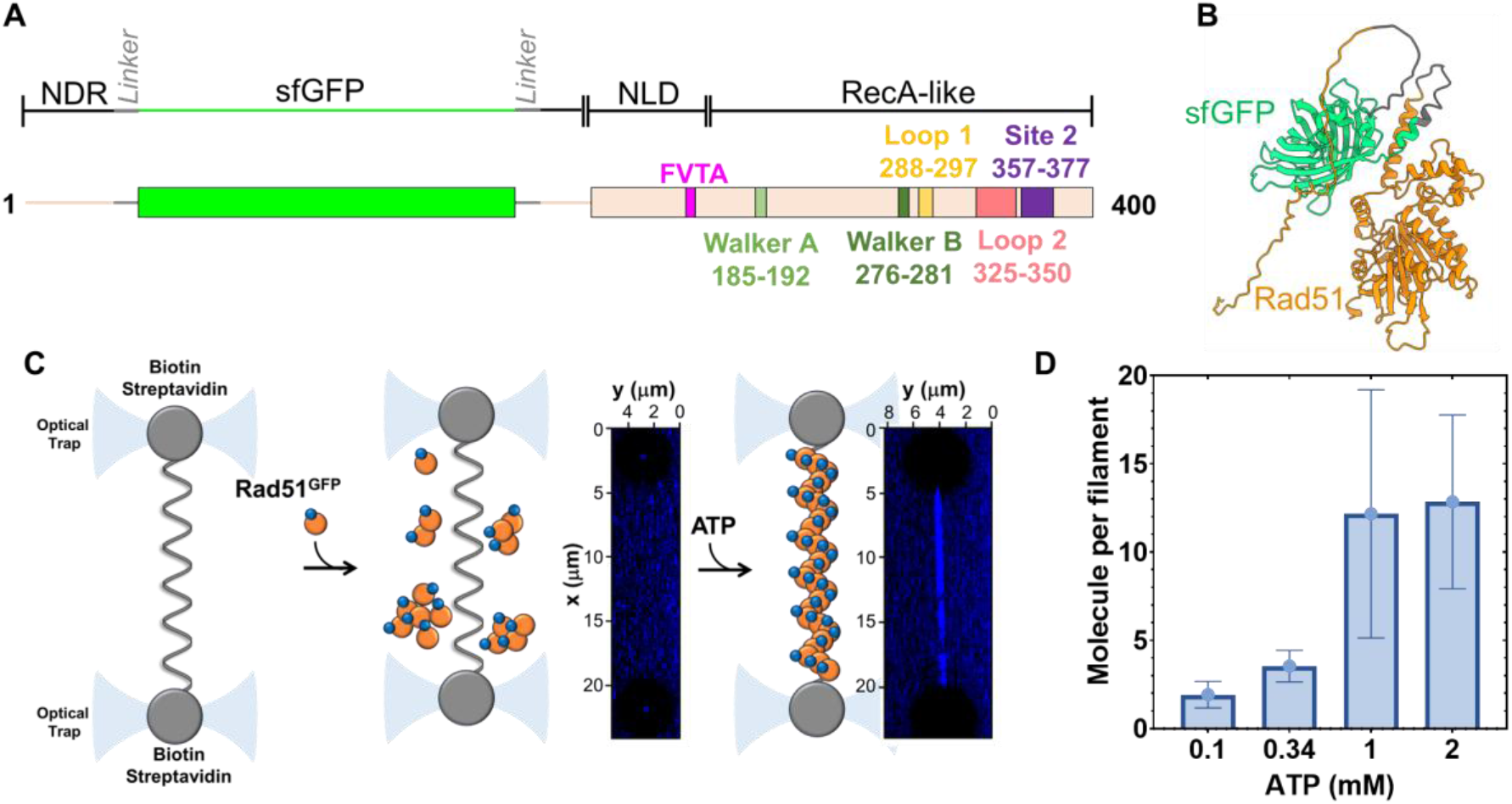
Direct visualization of Rad51 binding events on single-stranded DNA. **A)** Schematic of the sfGFP-Rad51 construct. The sfGFP is engineered at position 55 in the non-conserved N-terminal region of *S. cerevisiae* Rad51. Two 16 aa flexible linkers flank the sfGFP and required for maintaining Rad51 function. **B)** An AF-model of the Rad51^GFP^ protein. **C)** Optical trap analysis of Rad51^GFP^ binding to ssDNA. No binding is observed in the absence of ATP. **D)** Rad51^GFP^ binding to ssDNA is dependent on the ATP concentration. Data from n=15 tethers is shown, and Std. dev of the data are plotted.

To investigate the mechanism of Rad51 nucleoprotein filament formation, we performed confocal fluorescence imaging of recombinantly purified sfGFP-Rad51 (Rad51^GFP^; **Supplementary Figure 5**) on ssDNA under mechanical tension using a LUMICKS C-Trap, an instrument that combines dual optical traps with confocal scanning fluorescence microscopy with single fluorophore sensitivity^40^. Long ssDNA was created by stretching lambda phage dsDNA (∼48.5 kbp) that was tethered to two streptavidin-coated beads using biotin molecules that were positioned on the same strand. One strand was mechanically stretched until the complementary strand fell off due to high tension^41,42^. Formation of ssDNA was confirmed by fitting to a Freely Jointed Chain Model (FJC). Rad51^GFP^ was then allowed to bind to the ssDNA under 5 pN tension for 10-20 seconds before imaging. A 2D scan of the designated imaging area was performed to locate the confocal plane of the bound protein, followed by a line-scan along the protein-bound region over time, creating a kymograph (**Figure 4C**). Rad51 filament formation experiments were carried out either in the channel containing Rad51^GFP^, allowing new binding events to occur over time, or by binding in the Rad51^GFP^ channel followed by imaging in a separate channel to reduce background noise. As expected, Rad51^GFP^ does not bind to ssDNA in the absence of ATP, but forms filaments in an ATP concentration dependent manner (**Figure 4C & D**). We observed clear binding events of Rad51^GFP^ with signals resembling multimers. To quantify the multimeric state of each filament, we compared the signal intensity of a monomer with the initial signal intensity of the multimer to calculate the number of molecules in the filament (**Supplementary Figure 6**). The signal of the multimer/filament decreased over time, likely due to a combination of photobleaching and Rad51 dissociation. Thus, the initial signal intensity was used as a proxy for establishing the starting size of the filament.

### Rad52 sorts Rad51 to enhance Rad51 filament formation on non-RPA bound ssDNA

To investigate how Rad52 promotes formation of Rad51 filaments, we measured Rad51^GFP^ binding to ssDNA when in complex with Rad52. Using the optical trap setup described above, we moved the tethered ssDNA into the channel containing Rad51^GFP^ in the absence or presence of Rad52, incubated for 20s, and then imaged and recorded the frequency of Rad51^GFP^ filament formation per ssDNA tether (**Figure 5A**). We found that the presence of Rad52 increased the number of Rad51 multimer binding events (**Figures 5A, B & G**). When these experiments were performed on an RPA-coated ssDNA, no Rad51^GFP^ binding was observed (**Figure 5D**). The Rad52^ΔC^ protein that lacks the mode-1 interaction motifs does not stimulate Rad51^GFP^ binding (**Figure 5E**). In fact, there is a slight suppression of Rad51^GFP^ binding (**Figure 5G**). Finally, we observe poor dsDNA binding activity for Rad51^GFP^ (**Figures 5F & G**). These data show that the sorting function of Rad52 promotes formation of Rad51 filaments.

**Figure 5.**
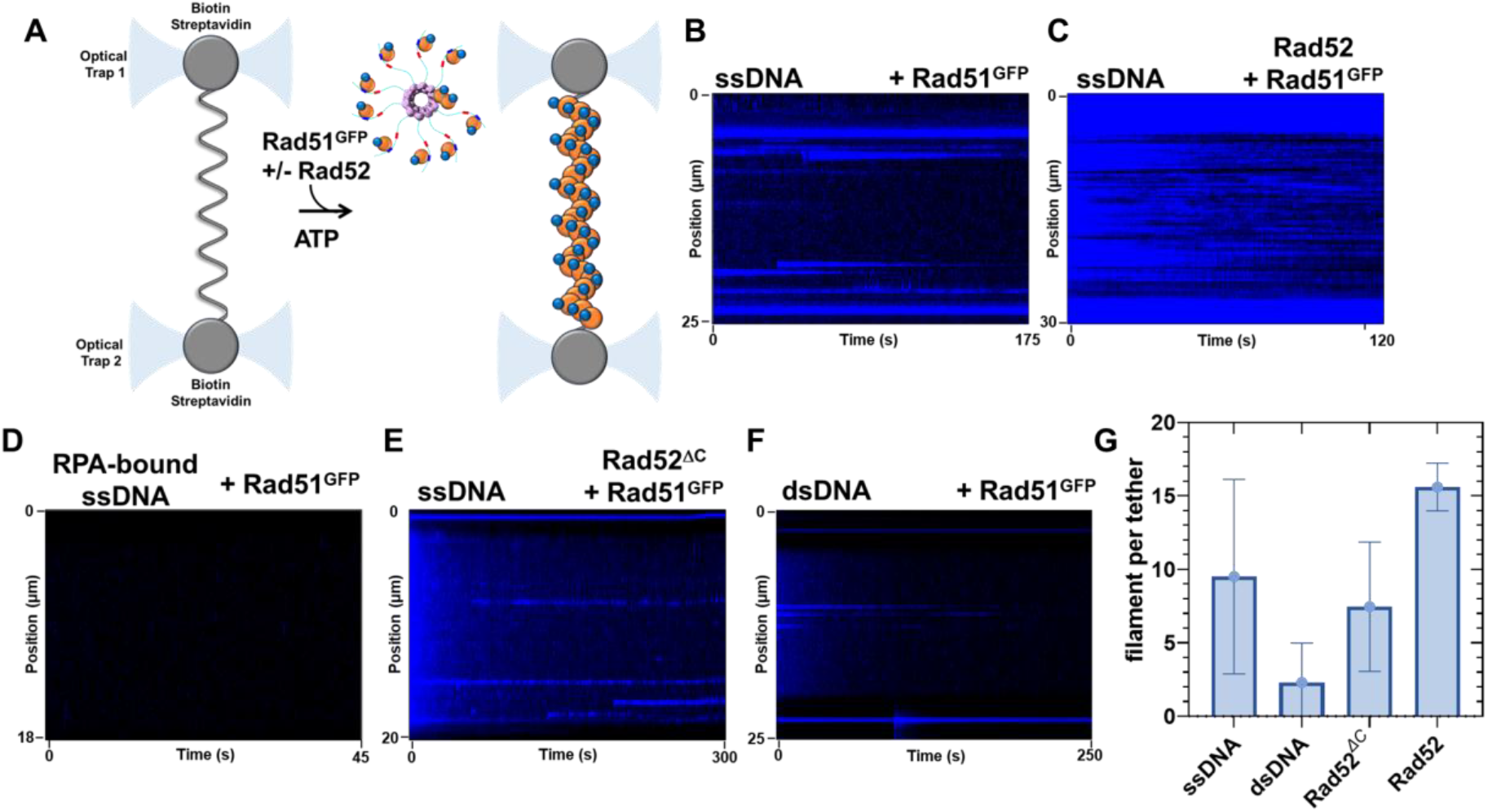
Rad52 promotes Rad51 binding to ssDNA. **A)** Schematic of the optical trap experiment showing Rad51^GFP^ binding to ssDNA in the absence/presence of Rad52. Kymographs of optical trap data in the **B)** absence or **C)** presence of Rad52 shows stimulation of Rad51^GFP^ binding to ssDNA. **D)** Kymograph showing no binding of Rad51^GFP^ to an RPA-coated ssDNA in the absence of Rad52. **E)** Rad52^ΔC^ that lacks the C-terminal Rad51 and RPA interaction domains poorly promotes Rad51^GFP^ binding to ssDNA. **F)** Poor binding of Rad51^GFP^ binding to dsDNA is observed. **G)** Quantitation of Rad51^GFP^ filaments formed under the denoted conditions. Representative data from a minimum of n=5 tethers per condition is shown, and Std. Dev. of the mean is plotted.

### Rad52 preferentially stacks Rad51 at the ss-dsDNA junction on RPA-coated ssDNA

The DNA used in the optical trap experiments described above is a ∼48 knt ssDNA substrate. However, the resected DNA substrate during HR pre-synapsis possesses both a ss-dsDNA junction and a 3’s free end. Technical needs of the optical trap to tether the DNA between two beads prevent us from being able to use a DNA with a free 3′ end. Thus, we utilized a modified gapped substrate that when stretched produces a ∼5 knt ssDNA gap in the middle (**Figure 6A**). In addition, an internal ATTO-647N fluorophore is positioned within the dsDNA region to infer polarity of the DNA and the two junctions. Here, we initiated the reaction by first coating the ssDNA with fluorescent RPA (site-specifically labeled with MB543 on DNA binding domain D; RPA^MB543^)^28,43,44^ in one channel and then moving the RPA^MB543^-ssDNA substrate to the next channel containing the Rad52-Rad51^GFP^ complex. In the first series of experiments, we maintained stoichiometric amounts of Rad52 (10 nM decamer) and Rad51^GFP^ (100 nM). The Rad52-Rad51^GFP^ complex bound to the ss-dsDNA junction with a clear preference for the junction (**Figure 6B**). In all our optical trap experiments, poor binding of Rad51^GFP^ to the dsDNA handles was observed.

**Figure 6.**
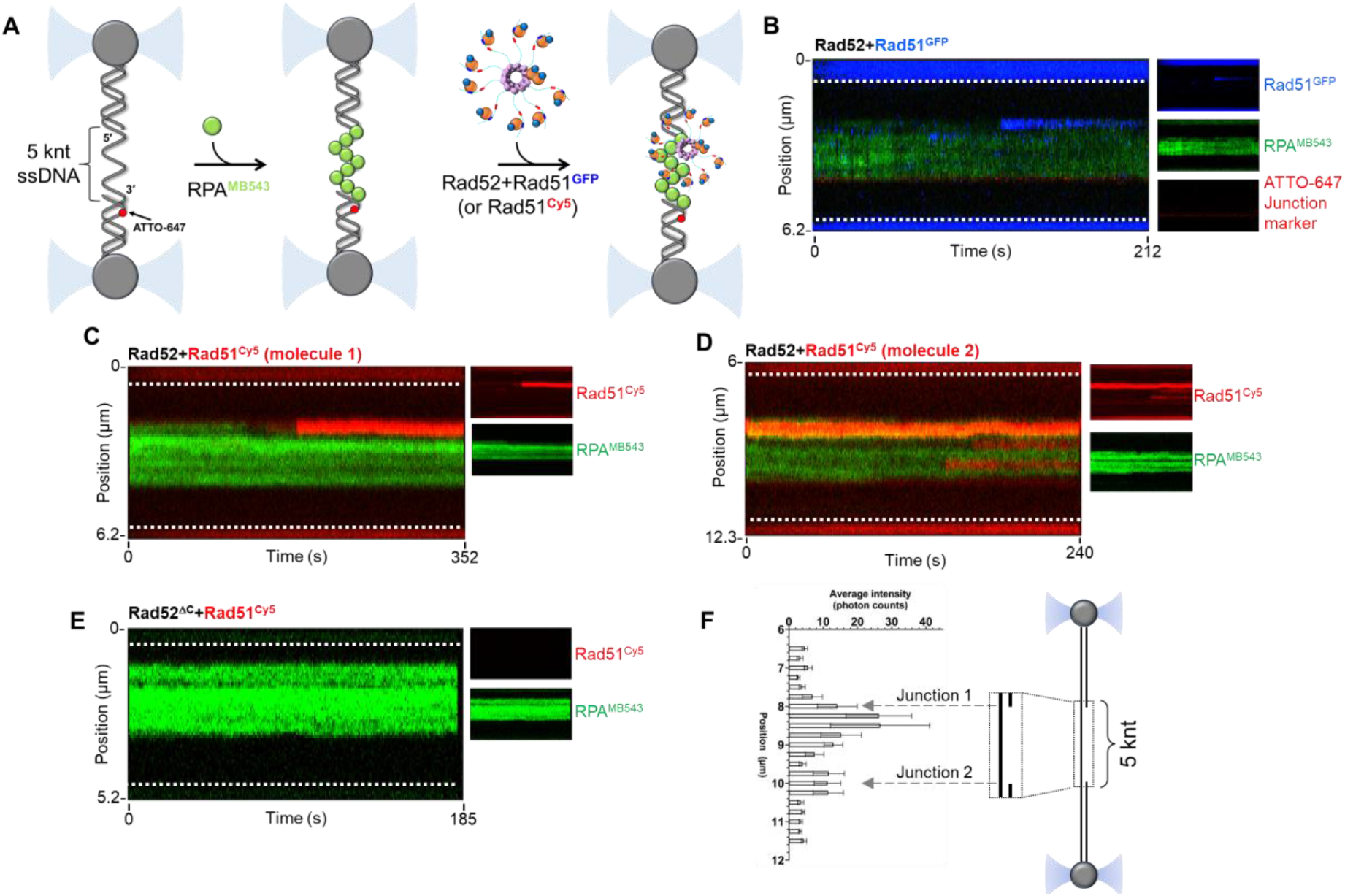
Rad51 binding to RPA-coated ssDNA in the presence of Rad52. **A)** Schematic of the optical trap experiment where the DNA is first preincubated with MB543-labeled-RPA (0.25 nM) and then moved to a channel containing fluorescent Rad51 (50 nM) in the presence of Rad52 or Rad52^ΔC^ (5 nM decamer). **B)** Stacks of frames (kymograph) were recorded by continuous confocal scanning along the DNA axis. The kymograph shows the binding of Rad51^GFP^ (blue) to the RPA^MB543^-coated ssDNA gap (green) in the presence of Rad52 and 5 mM ATP. Insets on the right side are single-color representations of the same kymograph. The Rad52-Rad51^GFP^ complex shows preferential binding to a position close to the junction. **C & D)** Kymographs of two DNA molecules showing the binding of Rad51^Cy5^ (red) to the RPA^MB543^-coated ssDNA gap (green) in the presence of Rad52 and 5 mM ATP. The Rad52:Rad51^Cy5^ complex also shows preferential binding to a position close to the junction. Rad51^Cy5^ binding to internal parts of the ssDNA gap, although of weaker intensity, can be also observed. Insets on the right side are single-color representations of the same kymograph. **E)** Kymograph recorded in the presence of Rad51^Cy5^, Rad52^ΔC^, and ATP. No binding of the Rad52^ΔC^-Rad51^Cy5^ complex is observed. The white dotted lines denote the boundaries of the optical traps/beads. **F)** Rad51^Cy5^ fluorescence signal intensity profile across DNA tethers averaged over all molecules analyzed (n = 8). Locations of the two ss-dsDNA junctions are indicated by gray dashed arrows. Preferential localization of the Rad51^Cy5^ signal adjacent to the junction is observed. Std. Err. of the mean is plotted.

The Rad51^GFP^ protein is fully functional in supporting Rad51 filament formation *in vivo*^29^. *In vitro*, Rad51^GFP^ interacts with Rad52 similar to untagged Rad51 (**Supplementary Figure 7**). To be certain that our observations do not arise due to some unforeseen artifacts of the linkers and/or the sfGFP, we generated an alternate fluorescent version of Rad51 by engineering a ybbR tag^45^ at position 55 in the same position as sfGFP and without the use of the two additional 16 amino acid linkers. A Cy5 fluorophore was attached to the ybbR tag using 4’
s-phosphopantetheinyl transferase (Sfp synthase) and coenzyme-A conjugated Cy5 (CoA-Cy5)^45^. The Rad51^GFP^ and Rad51^ybbR^ proteins display similar Rad52 binding properties in mass photometry analysis (**Supplementary Figure 7**). In optical trap experiments, the Rad51^Cy5^-Rad52 complex also preferentially binds adjacent to the ss-dsDNA junctions (**Figures 6C, D & F**). The Rad51^Cy5^ fluorescence signal was brighter and more photostable and thus used for downstream optical trap experiments. In addition, the ybbR construct allows us to vary the fluorophore on Rad51 as needed for multi-color single-molecule and ensemble experiments. Finally, tested whether a Rad52^ΔC^-Rad51^Cy5^ complex binds to the RPA^MB543^-coated ssDNA. No signal for Rad51^Cy5^ binding was observed when bound to Rad52^ΔC^ suggesting that the C-terminus of Rad52 is required for junction localization. The C-terminus of Rad52 possesses an RPA-binding motif in addition to the Rad51 mode-1 interaction and sorting functions uncovered in this study. Rad52^ΔC^ does not interact with RPA in AUC^SV^ analysis (**Supplementary Figure 8**). Thus, the defect in Rad51 binding we observe might arise from a lack of interaction of Rad52^ΔC^ with RPA, and/or loss of Rad51 sorting.

## Discussion

The formation of Rad51 nucleoprotein filaments during pre-synapsis is an essential step to drive HR and is promoted by mediator proteins such as Rad52 in yeast and BRCA2 in humans^4^. Since RPA binds to ssDNA with higher affinity compared to Rad51, mediator proteins function to overcome this thermodynamic barrier. Elemental to this process are physical interactions of the mediator protein with RPA, Rad51, and the DNA. However, knowledge of how these binding properties and assembly of the protein complexes enact Rad51 filament formation are poorly resolved. Here, we uncover several functional features of the yeast Rad52 mediator protein that showcase functional similarities to human BRCA2.

Rad52 is a homodecamer and each subunit possesses RPA and Rad51 binding motifs^21^. Our data shows that Rad51 interactions are driven by two distinct binding modes. Mode-1 is enacted by a Rad51 binding site in the disordered C-terminal region of Rad52 whereas mode-2 is driven through Rad51 interactions with the N-terminal ordered ring structure of Rad52. We here show that mode-1 interactions shift Rad51 from a polydisperse complex to monomers. This function resembles the Rad51-monomerization properties of the BRC-repeats of BRCA2^36,46,47^. In structural studies, mode-2 interactions are enacted in an asymmetric manner with Rad51 molecules stacked alongside one Rad52 subunit in the ring^21^. Thus, we propose a ‘*sort and stack*’ mechanism for Rad52-induced preparation/processing of Rad51 towards filament formation (**Figure 7**). Mediator proteins such as BRCA2 have been suggested to sequester recombinases such as RAD51 as a mechanism to keep them inactive and to suppress spurious interactions with DNA and prevent inhibition of other DNA metabolic processes^48^. Similarly, in prokaryotes, stacks of RecA oligomers were observed and to form dead-end structures and it was proposed that these must first disassemble in smaller oligomeric forms prior to active filament formation and HR^49^. Whether Rad52 serves such protective roles in yeast remains to be established.

**Figure 7.**
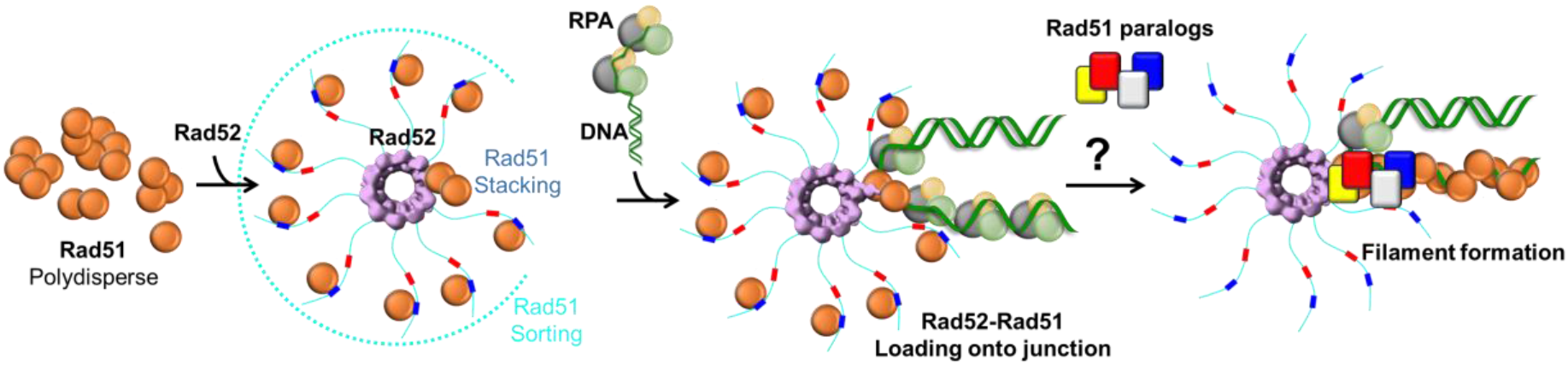
A sort and stack model for mediator-promoted Rad51 filament formation on RPA-coated ssDNA during HR. Rad51 is polydisperse in solution and is sorted into defined monomeric units by Rad52. There are two modes of interaction between Rad52 and Rad51. Mode-1 sorts Rad51 into monomers whereas mode-2 is asymmetric and occurs at one position in the Rad52 ring. The Rad52-Rad51 complex engages the RPA-bound resected ssDNA during pre-synapsis and preferential binding to the ss-dsDNA junction is observed. However, Rad52 is not sufficient to promote Rad51 filament growth. Whether Rad51-paralogs drive this transition into filament growth remains to be uncovered and its role/binding position in this model is speculative.

Our studies reveal a functional relevance for the sort and stack mechanism as Rad52-Rad51 complexes are preferentially loaded in the vicinity of the ss-dsDNA junction. In the context of pre-synapsis, there will be only one junction available owing to resection of the DSB and resulting in a 3′ ssDNA overhand. Thus, the Rad52-Rad51 complex will bind to this region and filament formation and growth will subsequently proceed towards the 3’
s end of the resected ssDNA. The junction preference and Rad52-Rad51 binding is lost when the C-terminus of Rad52 is deleted. This could either arise from the inability of Rad52^ΔC^ to either sort Rad51 and/or from a loss of RPA interactions. An intriguing observation is that the Rad52-Rad51 complex binds to the RPA-coated ssDNA in the absence or presence of ATP. Since Rad51 binds to DNA only in the presence of ATP, and neither RPA nor Rad52 possess dsDNA binding activity, the engagement of this complex at the junction is likely driven by physical interactions between RPA and Rad52. An alternative possibility is that either Rad52 or RPA possess DNA melting activity at the junction and the melted DNA structure provides a preferred binding location for the Rad52-Rad51 complex. These processes are likely further complicated in the presence of chaperone proteins such as Rtt105 that can influence both RPA and Rad51 interactions with ssDNA^42,50-53^. For example, in the absence of ssDNA, Rtt105 sequesters RPA and blocks physical interactions with Rad52^42^. In the presence of ssDNA, Rtt105 is remodeled to release the domains of RPA thereby promoting Rad52 binding. In addition, Rad52 engages the ssDNA-RPA complex by selectively remodeling DNA binding domain-D^43^, which would reside away from the junction due to the defined 5′->3′ binding polarity of the domains of RPA^54^. While the optical trap data presented here gives a bird’s eye view of the binding events at the junction, details of the underlying mechanisms are certainly more complex and need to be deciphered.

In addition, in the presence of ATP, we do not see rapid directional growth of the Rad51 filament on RPA-coated ssDNA. This property does not change even when excess Rad51 is introduced in the reaction. There are two possible explanations: a) Rad52 is actively suppressing filament growth, or b) other factors such as Rad51-paralogs are required for Rad51 filament formation after the initial nucleation of the Rad52-Rad51 complex (**Figure 7**). In addition to the lack of Rad51 filament growth, we observe no binding of Rad51 to dsDNA when Rad52 is present in the reaction. Thus, during pre-synapsis, Rad52 functions to actively suppress Rad51-dsDNA interactions while guiding localization of Rad51 to the junction. How the sorted Rad51 molecules are passed on from Rad52 onto the DNA remains another open question that needs to be investigated.

While yeast Rad52 and human BRCA2 are functional homologs, they share no structural similarity. However, the sort and stack mechanism that we describe here for Rad52 share many similarities to BRCA2. BRCA2 interacts with RAD51 through eight BRC repeats (1-8)^46,55^. The BRC repeats promote RAD51 binding to ssDNA while suppressing dsDNA interactions^36,55-59^. BRC1 and BRC4 have been shown to shift RAD51 to monomers^36^. Finally, a TR2 region in the C-terminus of BRCA2 interacts with filamentous RAD51 on DNA^47^. Thus, the BRC repeats could be envisioned as the sorter while TR2 region could be the stacker for pre-synaptic RAD51 filament formation. Optical trap studies of BRCA2 also show preferential localization towards the ss-dsDNA junction on RPA-coated DNA^56,58,60^. In addition, no growth of the RAD51 filament is observed in these BRCA2 optical trap studies. Thus, it is reasonable to assume that mediator proteins alone are not sufficient to facilitate filament growth, but function to localize a high concentration of monomeric Rad51 molecules in the vicinity of the ss-dsDNA junction. These studies now enable us to arrive at a reasonable consensus as to how yeast and human mediator proteins function to promote pre-synapsis during HR.

## Acknowledgements

This work was supported by grants from the National Institutes of Health NIGMS R35GM149320 to E.A., R35Gm122569 to T.H. E.A. We also acknowledge instrumentation support from the NIH Office of the Director S10OD030343 to E.A., and S10OD28650 to B.B. Funding for the Montana State Mass Spectrometry Facility was made possible in part by the MJ Murdock Charitable Trust, NIGMS P20GM103474, and the MSU Office of Research and Economic Development.

## Author Contributions

J.D. – performed AUC and MP experiments, generated Rad52 constructs, performed experiments, and generated figures. Assisted with manuscript preparation.

A.M. – performed MP experiments, generated sfGFP-Rad51 and Rad51^Cy5^ constructs and proteins, performed experiments, generates figures. Assisted with manuscript preparation.

S.P. and T.H. – performed optical trap experiments and analysis of Rad51 interactions on ssDNA. Manuscript edits and revisions.

M.G. and A.M.– performed optical trap experiments on the gapped DNA substrate. Manuscript preparation.

V.K. – generated fluorescent proteins for the study.

A.T. – provided information and advice on design of the sfGFP-Rad51 construct. Manuscript revisions.

B.B. and M.T. – performed XL-MS experiments and analysis. Manuscript edits and revisions.

E.A. – Study design and directed research. Manuscript generation.

## Methods

### Plasmids

sfGFP-Rad51 and yBBR-Rad51 were synthesized as codon-optimized ORFs and engineered into RSF-Duet1 plasmids (**Supplementary Figure 9**; Genscript Inc.). The sfGFP-Rad51 construct was designed as described^29^ and the sfGFP was engineered with flanking 16 aa linkers at the 55^th^ amino acid of the Rad51 coding region. The ybbR tag^45^ was also engineered at the 55^th^ position in the Rad51 coding region, but no additional linkers were required to generate active protein. Please see the Supplementary Information for amino acid composition of both constructs. The *S. cerevisiae* Rad52-pTXB1 plasmid (New England Biolabs Inc.) coding for an in-frame C-terminal chitin binding domain (CBD) tag was a kind gift from Dr. Eric Greene (Columbia University). Rad52^*ΔC*^, Rad52^*ΔN*^, and Rad52^*ΔN\**^ were generated as described^21^.

### Proteins

*Saccharomyces cerevisiae* Rad51, sfGFP-Rad51, and ybbR-Rad51 proteins were recombinantly overproduced using BL21(DE3)pLysS cells and purified using a previously described method^8^. A single transformant was grown in LB broth supplemented with 50 µg/mL kanamycin at 37ºC to an OD_600_ of 0.5-0.7. Protein overexpression was induced by the addition of 0.4 mM IPTG at 18ºC. After overnight induction at 18ºC, the cells were harvested and suspended in resuspension buffer (100 mM Tris-Cl, pH 8.0, 5 mM EDTA pH 8.0, 1 M NaCl, 1 M Urea, 5 mM beta-mercaptoethanol (β-ME), 10% w/v sucrose, 1X protease inhibitor cocktail, and 10% v/v glycerol). All downstream steps were carried out at 4ºC. Cells were first lysed with 0.4 mg/ml lysozyme for 30 min and sonicated for a total of 2 minutes in batches of 30-second ON-OFF cycles. The lysate was spun at 38,772 xg for 60 min and the clarified lysate was subjected to 40% w/v ammonium sulfate precipitation (0.24 g/mL) on ice. The solution was subsequently spun at 12,298 xg for 60 min. The pellet was resuspended in buffer Q^0^ (20 mM Tris-Cl, pH 7.5, 1 M Urea, 10% v/v glycerol, 0.5 mM EDTA, pH 8.0, 1 mM β-ME, and 1X protease inhibitor cocktail. The resuspended lysate was fractionated over a fast-flow Q-Sepharose column (Cytiva Inc.) and eluted with Q^100^-Q^700^ gradient (superscripts denote the NaCl concentration [mM] in the respective buffers). Fractions containing Rad51 were pooled and diluted with H^0^ buffer (30 mM Tris-Cl, pH 7.5, 0.5 mM EDTA pH 8.0, 0.5 mM β-ME, and 10% v/v glycerol) to match the conductivity value of a H^100^ buffer and fractionated over a fast-flow Heparin column (Cytiva Inc.) equilibrated with buffer H^100^. Rad51 was eluted with a linear gradient of H^100^-H^1000^. Heparin fractions containing Rad51 were concentrated to ∼5 mL using an Amicon Ultra spin concentrator (Sigma Inc.). The concentrated Rad51 was loaded onto a Hi-Load 16/600 Superdex 200 pg size exclusion column (Cytiva Inc.) equilibrated with buffer (20 mM Tris-Cl, pH 7.5, 0.5 mM EDTA pH 8.0, 1 mM β-ME, 10% v/v glycerol, and 100 mM NaCl). Fractions containing Rad51 were pooled and dialyzed against storage buffer (20 mM Tris-Cl, pH 7.5, 0.5 mM EDTA, pH 8.0, 1 mM β-ME, 20% v/v glycerol, and 100 mM NaCl), flash frozen using liquid nitrogen, and stored at -80°C. Protein concentrations were measured using the following extinction coefficients ε_280_: 11,920 M^-1^ cm^-1^ for Rad51 and ybbR-tagged Rad51, and 33,810 M^-1^ cm^-1^ for sfGFP-Rad51.

Sfp-synthase was overproduced using a pBAD-MBP-His-SFP plasmid (a kind gift from New England Biolabs - Addgene #141141). The plasmid was transformed into BL21(DE3)pLysS cells and transformants were grown in LB broth supplemented with 50 µg/mL ampicillin at 37°C to an OD_600_ of 0.6. Protein overexpression was induced by the addition of 0.2% (v/v) L-arabinose. After overnight induction at 16°C, the cells were harvested and suspended in resuspension buffer (500 mM NaCl, 5 mM imidazole, and 20 mM Tris-Cl, pH 8.0). All downstream steps were carried out at 4°C. Cells were lysed with 0.4 mg/ml lysozyme and sonicated for a total of 2 minutes in batches of 30-second ON-OFF cycles. The lysate was spun at 38,772 xg for 60 min. The clarified lysate was first batch-bound to 5 ml Ni-NTA resin (Gold Biotechnology Inc.) for 3 hours. The beads were subsequently washed with resuspension buffer, followed by stepwise elution using resuspension buffer containing 100, 200, or 400 mM imidazole, respectively. The eluate containing Sfp-synthase was then concentrated to ∼5 mL using an Amicon Ultra spin concentrator and fractionated over a Hi-Load 16/600 Superdex 200 pg size exclusion column equilibrated with S200 buffer (20 mM Tris-Cl, pH 8.0, 500 mM NaCl, 10% v/v glycerol, and 1 mM β-ME). Please note that Sfp-synthase elutes as two distinct fractions on SEC. While both fractions contain clean protein, the lower molecular mass peak is catalytically active and used for labeling reactions described in this study. Fractions containing Sfp-synthase were flash frozen using liquid nitrogen, and stored at -80°C. Sfp-synthase concentration was measured spectroscopically using ε_280_ =28,880 M^-1^ cm^-1^.

*Saccharomyces cerevisiae* Rad52, Rad52^*ΔC*^, Rad52^*ΔN*^, and Rad52^*ΔN\**^ proteins were purified as described^21^. Unlabeled and MB543-labeled *Saccharomyces cerevisiae* RPA were purified as described^28,43,44,61^. For this study, RPA labeled in DNA binding domain D^44^ was used in the optical trap experiments.

### Rad51 labeling with CoA-Cy5

The CoA-Cy5 conjugate was synthesized as described^45^. CoA-Cy5 was generated by preparing a 1000 µL reaction by adding 1.6 mg CoA-trilithium salt (Sigma Inc.) in 750 µL 100 mM Na_3_PO_4_ pH 7.0 to a solution containing 1 mg Cy5-maleimide dye in 250 µL DMSO. This reaction mixture was stirred at room temperate in the dark for 60 minutes. The mixture was then passed through a C18 reversed-phase HPLC column (Avantor-VWR Inc.). Peak fractions carrying the CoA-Cy5 conjugate were collected and stored at -20°C. For the labeling reaction, 5 µM CoA-Cy5, 5 µM ybbR-Rad51, and 0.1 µM SFP synthase were mixed in a 500 µl reaction in labeling buffer (50 mM HEPES, pH 7.5, and 10 mM MgCl_2_). The reaction was incubated at room temperature for 30 minutes in the dark and then passed through a Biogel P-4 column (Bio-Rad Inc.) equilibrated with buffer (20 mM Tris-Cl, pH 8.0, 500 mM NaCl, and 10% v/v glycerol). The final protein concentration of Rad51-Cy5 and the labeling efficiency were calculated with the ε_280_ =11,920 M^-1^ cm^-1^ for the protein and ε_650_= 250,000 M^-1^ cm^-1^ for Cy5. In addition, the value at 280 was adjusted using a correction factor of 0.03 to account for contribution of the Cy5 to the absorption signal. The Rad51-Cy5 protein was flash-frozen using liquid nitrogen and stored at - 80°C. During experiments, the Rad51-Cy5 protein was not subject to repeated freeze-thaw cycles.

### Crosslinking Mass spectrometry (XL-MS)

XL-MS experiments and analyses were carried out as described^21^. Briefly, stock solutions of Rad52 and Rad51 were diluted to 1.8 mg/mL and 2.2 mg/mL, respectively in buffer (30 mM HEPES, 200 mM KCl, pH 7.8) and incubated together for 30 min. Crosslinking reaction and sample processing were conducted in similar fashion as described before (ref21). Briefly, stock solutions of Rad52^ΔC^ and Rad51 were diluted to 0.9 mg/mL and 1.5 mg/mL, respectively in buffer (30 mM HEPES, 300 mM NaCl, pH 7.8) and incubated together (Rad52^ΔC^ to Rad51 molar ratio as 1 : 10) for 5 min on ice. The diluted proteins were incubated with primary amine reactive 3 mM (final) bis(sulphosuccinimidyl)suberate (Sigma) and 20 µL of the sample was taken at various time points (0, 15, and 30 minutes) and immediately quenched with 2 µL of 1 M ammonium acetate (Sigma). Quenched samples were diluted with 4X Laemmli gel loading buffer to a final volume of 30 µL, boiled for 5 min and then resolved on 4-20% (w/v) gradient SDS-PAGE gels (Bio-Rad) with Tris-glycine buffer. To visualize protein, gels were stained with Gelcode blue safe protein stain (ThermoFisher Inc.). The major protein bands from 15 min time point were excised, briefly destained, and proteins were reduced with 10 mM dithiothreitol (Sigma) for 30 min at 56 °C and then alkylated with 50 mM iodoacetamide (Sigma) for 25 min at room temperature in a dark. Next, proteins were digested with 100 ng porcine sequencing grade modified trypsin (Promega) overnight at 37 °C. LCMS experiments were performed as described before^62^. Briefly, tryptic peptides were separated by reverse phase XSelect CSH C18 2.5 um resin (Waters) on an in-line 150 × 0.075 mm column using an UltiMate 3000 RSLCnano system (ThermoFisher Inc.). Peptides were eluted using a 60 min gradient from 98:2 to 40:60 solvent A:B ratio (solvent A = 0.1% formic acid, 0.5% acetonitrile; solvent B = 0.1% formic acid, 99.9% acetonitrile). Eluted peptides were ionized by electrospray (2.4 kV) followed by mass spectrometric analysis on an Orbitrap Eclipse Tribrid mass spectrometer (Thermo). MS data were acquired using the FTMS analyzer in profile mode at a resolution of 120,000 over a range of 375 to 1400 m/z with advanced peak determination. Following HCD activation, MS/MS data were acquired using the FTMS analyzer in profile mode at a resolution of 15,000 over a range of 150 to 2000 m/z with a stepped collision energy of 27-33%. Raw data files were converted to mgf format using ProteoWizard 3.0^63^ and then uploaded to Spectrum Identification Machine 1.5.6^64^ for crosslinks identification. Crosslinking patterns between bound and unbound proteins were compared across the same migration range in the gel, monomers to monomers, dimers to dimers, etc.

### Mass photometry

Mass photometry measurements were carried out on a TwoMP instrument (Refeyn Inc.), which was allowed to warm-up for an hour prior to the experiments. Glass coverslips (No. 1.5H thickness, 24 × 50 mm, VWR Inc.) were cleaned by sequential sonication in isopropanol and deionized water for 15 minutes, respectively. Cleaned cover slips were dried under a stream of nitrogen. The molecular weight standard (β-amylase, with three species of 56, 112, and 224 kDa, respectively), Rad51, Rad52^ΔN^, and Rad52^ΔC^ were diluted in the same buffer comprised of 50 mM Tris-acetate, pH 7.5, 50 mM KCl, and 5 mM MgCl_2_. β-amylase was diluted to a final concentration of 200 nM, while Rad51, Rad52^ΔN^, and Rad52^ΔC^ were diluted to a final concentration of 100 nM. Three different stoichiometries of both Rad52^ΔN^ and Rad52^ΔC^ were tested separately (1:1, 3:1, and 5:1) with a fixed concentration of Rad51. For these experiments, 500 µL reactions were prepared in the same buffer with the relevant ratios of Rad52^ΔN^/Rad52^ΔC^ and Rad51 concentrations. For our experiments, we used a constant concentration of 100 nM for Rad51 against 100, 300, and 500 nM Rad52^ΔN^/Rad52^ΔC^. The reactions were spun down on a table-top centrifuge for one minute and then allowed to incubate at room temperature for precisely 20 minutes. The buffer was also maintained at room temperature throughout the analysis. For each measurement, a clean coverslip with a 6-well silicone gasket was placed onto the oil-immersion objective and samples were added into each well as described below. After aligning the laser approximately to the center of the first well, 15 µL of the buffer was added and focus was obtained. Then, 1 µL of the 200 nM β-amylase was added to the same well (final concentration of 12.5 nM) to obtain high-resolution data with significant number of particles (∼2500) over a 60 s recording. This data was used to perform mass calibration. The experimental samples were similarly added to their respective wells after finding focus with 15 µL of the buffer. For data analysis, single-particle landing events were identified and converted to mass units using the above-mentioned standard calibration, extracted from videos, and non-linear least squares fitted with Gaussian mixture model to quantify the underlying populations using the Refeyn DiscoverMP software.

### Mass photometry

Mass photometry measurements were carried out on a TwoMP instrument (Refeyn Inc.), which was allowed to warm-up for an hour prior to the experiments. Glass coverslips (No. 1.5H thickness, 24 × 50 mm, VWR Inc.) were cleaned by sequential sonication in isopropanol and deionized water for 15 minutes, respectively. Cleaned cover slips were dried under a stream of nitrogen. The molecular weight standard (β-amylase, with three species of 56, 112, and 224 kDa, respectively), Rad51, Rad52^ΔN^, and Rad52^ΔC^ were diluted in the same buffer comprised of 50 mM Tris-acetate, pH 7.5, 50 mM KCl, and 5 mM MgCl_2_. β-amylase was diluted to a final concentration of 200 nM, while Rad51, Rad52^ΔN^, and Rad52^ΔC^ were diluted to a final concentration of 100 nM. Three different stoichiometries of both Rad52^ΔN^ and Rad52^ΔC^ were tested separately (1:1, 3:1, and 5:1) with a fixed concentration of Rad51. For these experiments, 500 µL reactions were prepared in the same buffer with the relevant ratios of Rad52^ΔN^/Rad52^ΔC^ and Rad51 concentrations. For our experiments, we used a constant concentration of 100 nM for Rad51 against 100, 300, and 500 nM Rad52^ΔN^/Rad52^ΔC^. The reactions were spun down on a table-top centrifuge for one minute and then allowed to incubate at room temperature for precisely 20 minutes. The buffer was also maintained at room temperature throughout the analysis. For each measurement, a clean coverslip with a 6-well silicone gasket was placed onto the oil-immersion objective and samples were added into each well as described below. After aligning the laser approximately to the center of the first well, 15 µL of the buffer was added and focus was obtained. Then, 1 µL of the 200 nM β-amylase was added to the same well (final concentration of 12.5 nM) to obtain high-resolution data with significant number of particles (∼2500) over a 60 s recording. This data was used to perform mass calibration. The experimental samples were similarly added to their respective wells after finding focus with 15 µL of the buffer. For data analysis, single-particle landing events were identified and converted to mass units using the above-mentioned standard calibration, extracted from videos, and non-linear least squares fitted with Gaussian mixture model to quantify the underlying populations using the Refeyn DiscoverMP software.

### Analytical Ultracentrifugation

Sedimentation velocity experiments were performed with an optima analytical ultracentrifuge (Beckman-Coulter Inc.) using An-50Ti rotor at a speed of 40,000 rpm at 20°C. Samples were prepared by dialyzing against 30 mM HEPES, pH 7.8, 100 mM KCl, 10 % glycerol, 1 mM TCEP and concentrations of each component of samples are mentioned in the figure. Sample (380 µL) and buffer (400 µL) were filled in each sector of 2-sector charcoal quartz cells. Absorbance was monitored at 280 nm for samples using absorbance optics and scans were obtained at 3 min intervals. The density and viscosity of the buffer at 20°C was calculated using SEDNTERP^65^. The continuous distribution (c(s)) model was used to fit the AUC data by SEDFIT program^66,67^.

### Co-relative Optical Trap Experiment and analysis

A 48.5 kilo base (Kbp) Lambda DNA was prepared with three biotins on the 3′ and 5′ ends of the same strand. Lambda DNA was purchased from New England Biolabs (N3011). This linear DNA comes with 12 nucleotide (nt) 5′ overhangs. Three short oligos to anneal to the sticky ends were purchased from Integrated DNA Technologies Inc. Oligo 1: 5′-GGG CGG CGA CCT GGA CAA-3′; oligo 2: 5′-AGG TCG CCG CCC TTT TTT T/iBiodT/T /iBiodT/T/iBiodT/-3′; oligo 3: 5′-/iBiodT/T/iBiodT/ T/iBiodT/T TTT TTT AGA GTA CTG TAC GAT CTA GCA TCA ATC TTG TCC-3′. The DNA was stored in TE buffer (10 mM Tris-HCl, pH 8.0, and 0.1 mM EDTA). The final ssDNA construct was prepared as described^41^. Briefly, a Lumicks BV Inc. commercial optical trap with combined confocal microscopy was used. The system has a microfluidic setup with five experimental channels, each with a height of 100 µm. Re-usable glass slides provided by the company are extensively passivated before the experiment to create a uniform surface. Passivation was performed sequentially with 0.5 mL of each component in separate syringes: 0.1% BSA (Sigma Inc.) flowing at 1 bar for 5 minutes, then at 0.4 bar pressure for 25 minutes, followed by a 10-minute rinse with RNase-free water at 1 bar pressure for 5 minutes, and 0.4 bar pressure for 5 minutes. This was followed by 0.5% Pluronic F-127 which was flowed at 1 bar pressure for 5 minutes, then at 0.4 bar pressure for 25 minutes, followed by 1X Phosphate Saline Buffer (PBS) at 1 bar pressure for 5 minutes and 0.4 bar pressure for 5 minutes. Streptavidin-coated polystyrene particle beads of average size 4.8 µm [0.5% w/v] (Spherotech Inc.) were diluted 1:250 in 1X PBS containing 137 mM NaCl, 2.7 mM KCl, 8 mM Na_2_HPO_4_, and 2 mM KH_2_PO_4_ (10X PBS was purchased from Thermo Fisher Scientific USA Inc.). 100 ng of DNA was made in 0.5 mL of 1X PBS. DNA was captured between two streptavidin beads and mechanically denatured by moving one bead to the overstretched region to create ssDNA. ssDNA was confirmed by fitting the force-distance (FD) curve to the Freely Jointed Chain model (FJC) (contour length 48.5 kbp / 27.160 µm; persistence length 0.9 nm; stretch modulus 1000 pN) in real-time. DNA was held for 5 seconds in the fully ssDNA state, then returned to a required tension on the ssDNA position for the fluorescence experiments.

For characterization of Rad51 binding events on the long ssDNA experiments, 100 nM of sfGFP-Rad51 was used for each experiment. The buffer used was 100 mM NaCl, 50 mM HEPES pH 7.5, and 10 mM MgCl_2_. The ATP concentration was varied as denoted. The imaging buffer contained 0.8% (w/v) dextrose, 165 U/mL glucose oxidase, 2170 U/mL catalase, and 2-3 mM Trolox to increase the fluorescence lifetime of the fluorophores. Imaging settings were 0.1-0.5 ms exposure time (per pixel), with a constant pixel size of 100 nm. Excitation wavelengths were Green (575-625 nm) and Red (670-730 nm). Laser power was selected to maintain 5 micro-watt power at the objective. The data was analyzed by custom written python scripts. Pylake package of Lumicks was used in python to process the data. Scripts used can be accessed through the following Github link (https://github.com/spangeni/Rad51_eGFP-Paper). Further raw data can be made available upon request.

For characterization of Rad51 binding events on the gapped DNA substrates, data were collected on a different optical trap instrument with the following modifications to the procedure and analysis. Prior to use, the protein channels of the microfluidics chip were passivated according to Lumicks protocol with minor modifications. Briefly, 0.1% BSA resuspended in Rad51 buffer (50 mM Tris-Cl, pH 7.5, 50 mM NaCl, 10 mM MgCl_2_ and 0.2 mg/mL BSA) was flowed in for 20 min at 0.35 bar, followed by 0.5% Pluronic F-127 in the same buffer for 20 min at 0.35 bar. Finally, a minimum of 300 µL of Rad51 buffer flowed in to equilibrate the microfluidics chip. Biotinylated ds/ss DNA hybrid precursor (17.852 kbp, Cat # 00027Lumicks Inc.) and 4.35 µm Streptavidin-coated polystyrene beads from Lumicks were diluted in 1X running buffer (137 mM NaCl, 2.7 mM KCl, 1.12 mM phosphates, 5 mM sodium azide, and 0.5 mM EDTA, pH 7.4) according to manufacturer’s instructions. ds/ss DNA hybrid precursor was captured between two beads at 0.29 pN/nm trap stiffness, and ss DNA gap was generated by pulling the DNA in a low-salt buffer (0.1X running buffer). Formation of ssDNA gap was verified by comparison to worm-like chain model in the instrument-associated Bluelake software. In addition, ssDNA gap was confirmed by visualizing DNA using MB543-labeled yeast RPA protein which binds to only ssDNA with high affinity. Prior to visualization of the interaction of yeast Rad51/Rad52 with RPA-coated ssDNA, the three proteins were diluted in Rad51 buffer. The ss/dsDNA hybrid was stretched between two beads with force of 10 pN at a flow pressure of 0.1 bar. The DNA was moved to a microfluidics flow channel containing 0.25 nM RPA^MB543^ in imaging buffer (Rad51 buffer supplemented with 2 – 3 mM Trolox, 0.165 U/µL glucose oxidase, 2.17 U/ µL catalase and 0.8% (w/v) D-glucose monohydrate). After confirming the coating of the ssDNA gap with fluorescent RPA, the DNA was then moved to another microfluidics channel containing 50 nM or 100 nM Cy5-labeled Rad51 and 50 nM Rad52 or Rad52^DC^ in imaging buffer with or without 5 mM ATP. Confocal microscopy was carried out using green (561 nm) and red (638 nm) lasers, with pixel size of 100 nm and 2 ms per pixel. Stacks of frames (kymograph) recorded by continuous confocal scanning along the DNA axis over time were visualized using the Lumicks-developed Lakeview Pro software. Fluorescence intensity profiles of Rad51^Cy5^ along DNA were extracted from the kymographs from all molecules captured in the analysis for each condition. Intensity-position profiles were averaged, errors bars as standard errors of the means of fluorescence intensity at each position among all molecules were calculated and plotted in GraphPad Prism 10.3.0.

### Structural models

A model of the Rad51-Rad52 complex was generated by using the respective amino acid sequences as input for the web-service AlphaFold server (https://alphafoldserver.com/) powered by AlphaFold3^68^. Default settings were used to generate predicted structure models for Rad52-Rad51. Molecular graphics and analyses were performed using UCSF ChimeraX^69^. The top-ranked predictions are displayed in the figures.

