## Supplementary Information for "Rad52 sorts and stacks Rad51 at the DNA junction to promote homologous recombination"

**Supplementary Figures 1-9**

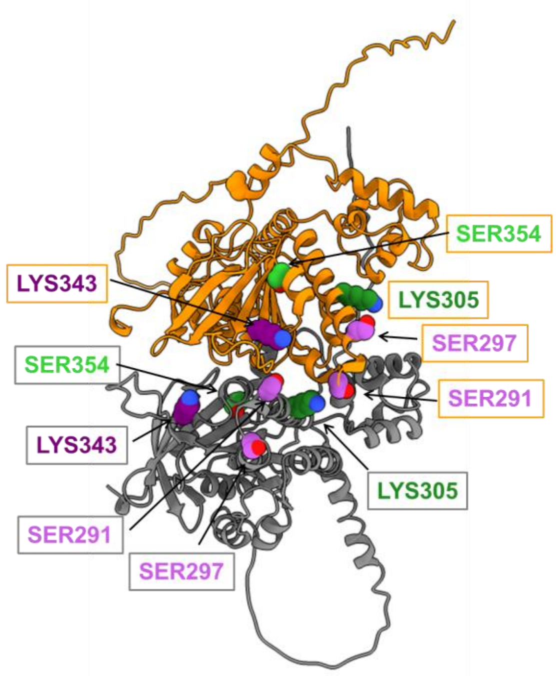


**Supplementary Figure 1. The intra/inter crosslinks in the Rad52-Rad51 complex map to the Rad51 oligomerization interface.** In crosslinking mass spectrometry analysis of the Rad2-Rad51 complex only three inter- or intra-crosslinks were identified within Rad51 (Figure 2C). When mapped onto the structure of an AF2 model of the yRad51 filament, these residues are situated within the oligomerization interface. Two Rad51 molecules in the filament are colored grey and orange. Lysine and serine residues are colored in green and purple, respectively.

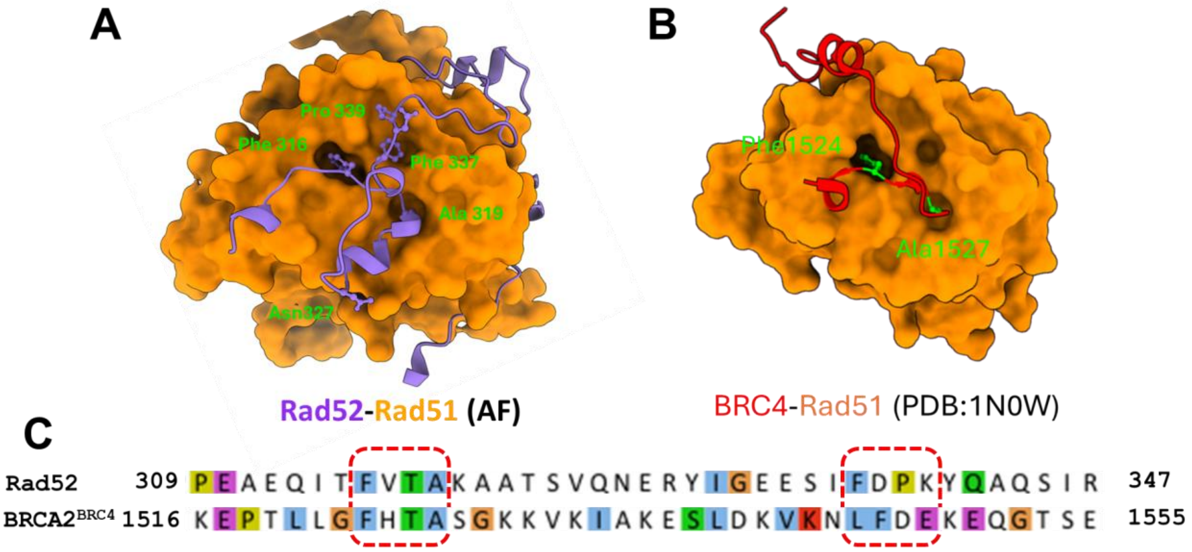


**Supplementary Figure 2. The Mode-1 Rad51 interaction features are conserved between yeast Rad52 and human BRCA2.** **A)** AlphaFold model of the C-terminus of Rad52 (purple) bound to a Rad51 monomer. Phe-316, Ala-319, and Phe-337 dock in to defined hydrophobic pockets in Rad51. **B)** Crystal structure of human RAD51 bound to the BRC4 peptide from BRCA2 (PDB:1N0W) is shown. Phe-1524 and Ala1527, that are homologous to Phe-316 and Ala-319 in the yeast Rad52 FVTA motif, dock into similar hydrophobic pockets. **C)** Sequence alignment of the Rad51 interaction regions between yeast Rad52 and human BRCA2. The FXTA and FDXK motifs are highlighted by the dotted red boxes.

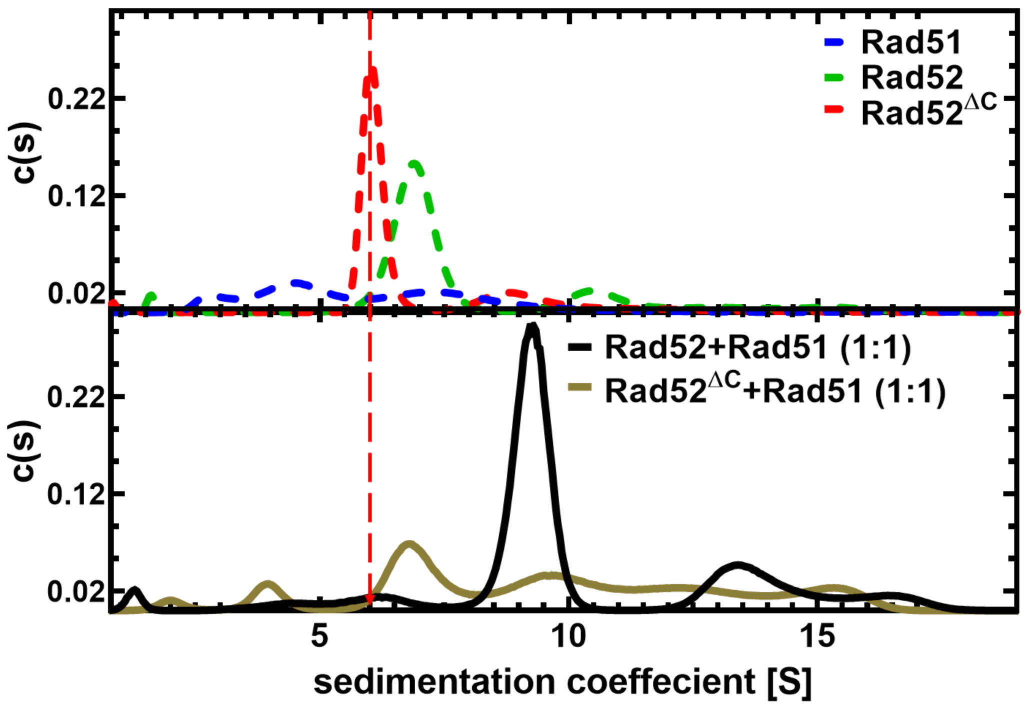


**Supplementary Figure 3. Sedimentation velocity AUC experiments show interactions between Rad52$^{\Delta C}$ and Rad51.** Top panel shows the sedimentation profiles of Rad51 (blue), Rad52 (red), and Rad52$^{\Delta C}$ (green). Bottom panel shows the sedimentation profiles of the Rad51-Rad52 (black) and Rad51-Rad52$^{\Delta C}$ (pale green) complexes. The Rad51-Rad52 complex is relatively uniform in comparison to the Rad51-Rad52$^{\Delta C}$ complex.

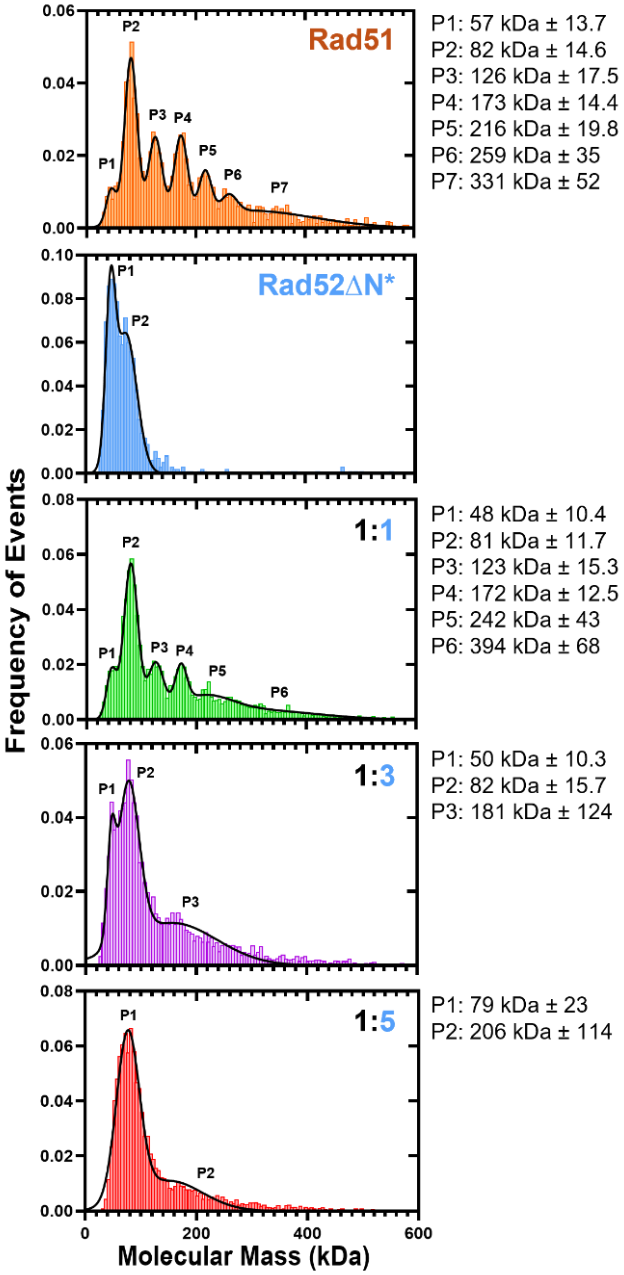


**Supplementary Figure 4. Mass photometry analysis of the Rad52$^{\Delta N*}$-Rad51 complex.** Increasing ratios of Rad52$^{\Delta N*}$ were premixed with Rad51 and analyzed using mass photometry. As the ratios are increased, the distribution of Rad51 in solution shifts from oligomers to monomers. The expected molecular weight of a Rad51 monomer is 43 kDa compared to 71 kDa for the Rad52$^{\Delta N*}$-Rad51 complex. The measured masses are denoted alongside each species observed in the experiment. Please note that the detection limit for accurate mass measurements in mass photometry is >30 kDa. Thus, the measured mass for the 19 kDa Rad52$^{\Delta N*}$ fragment is below the detection limit of the instrument.

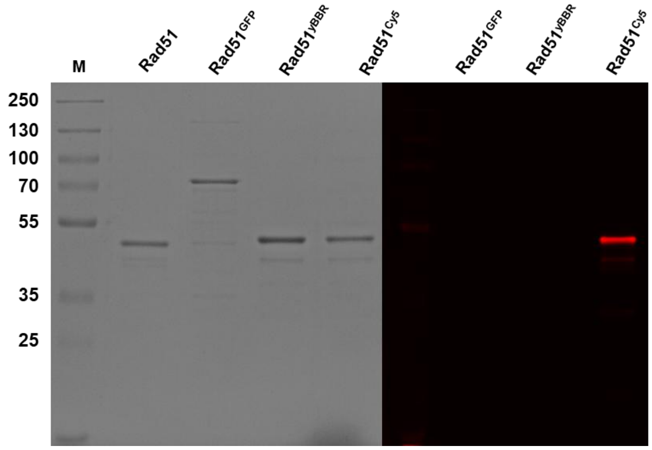


**Supplementary Figure 5. Generation of fluorescent Rad51 proteins.** SDS-PAGE analysis of the Rad51 and Rad51 variants are shown. Coomassie staining (left) and fluorescence imaging (right) of the same gel are shown.

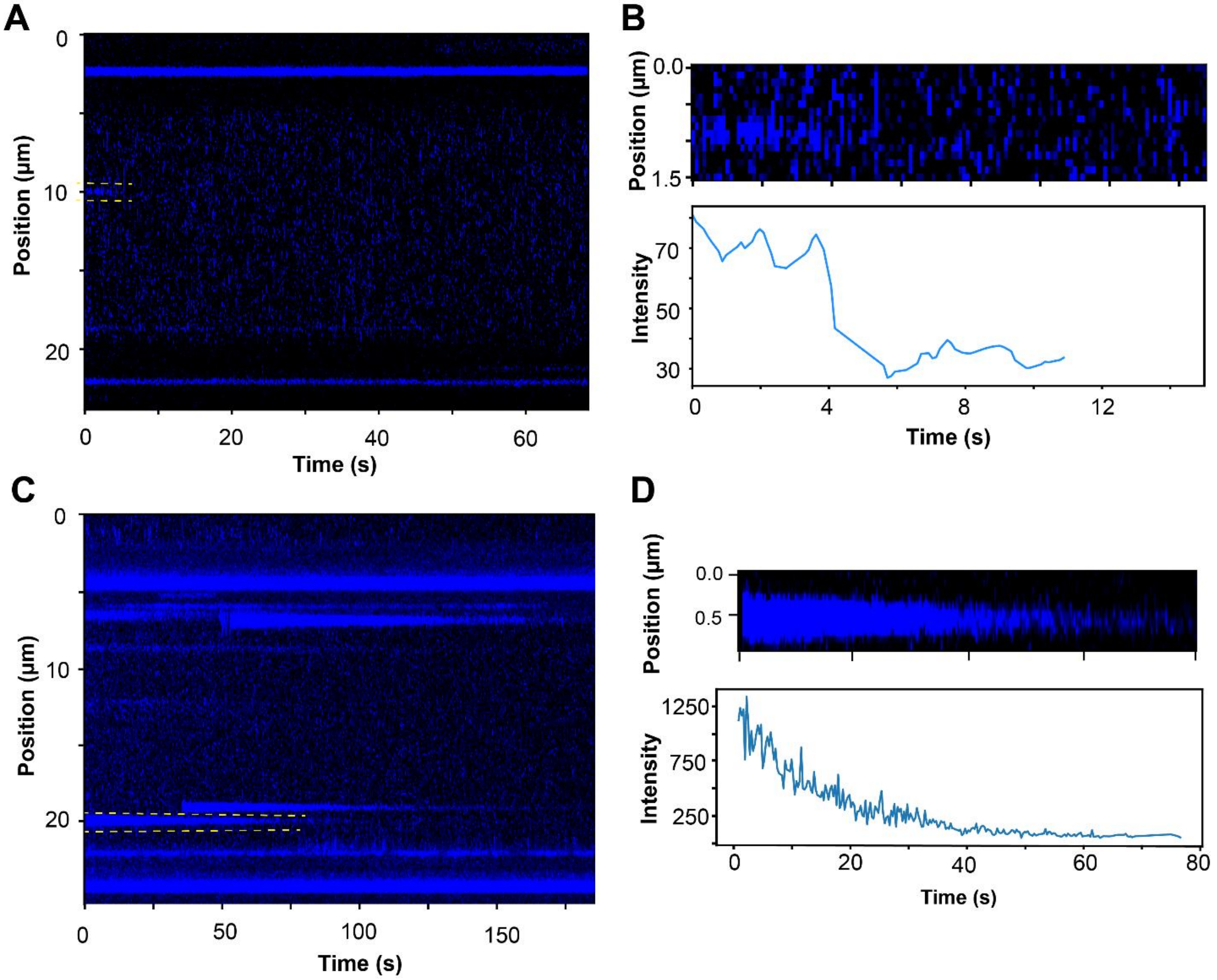


**Supplementary Figure 6. Analysis of optical trap measurements of Rad51$^{GFP}$ filaments on ssDNA.** Kymographs of Rad51$^{GFP}$ binding to ssDNA were recorded by **A)** transiently dipping the ssDNA in channel containing Rad51$^{GFP}$ or **D)** incubating the ssDNA for 20 sec to promote stable Rad51$^{GFP}$ binding. Yellow boxes are regions zoomed in for analysis in panels **B)** and **D)**, respectively. **B)** Shows transient Rad51 binding events. The intensity corresponds to smaller Rad51 binding units (monomer-trimer) and the dissociate rapidly. **D)** Rad51$^{GFP}$ signal that we observe upon longer incubation periods and interpret as filaments for our analysis. In these experiments on ssDNA, we see a stark time-dependent decrease in the Rad51$^{GFP}$ signal. This drop in intensity could be attributed to either dissociation of Rad51 and/or photobleaching.

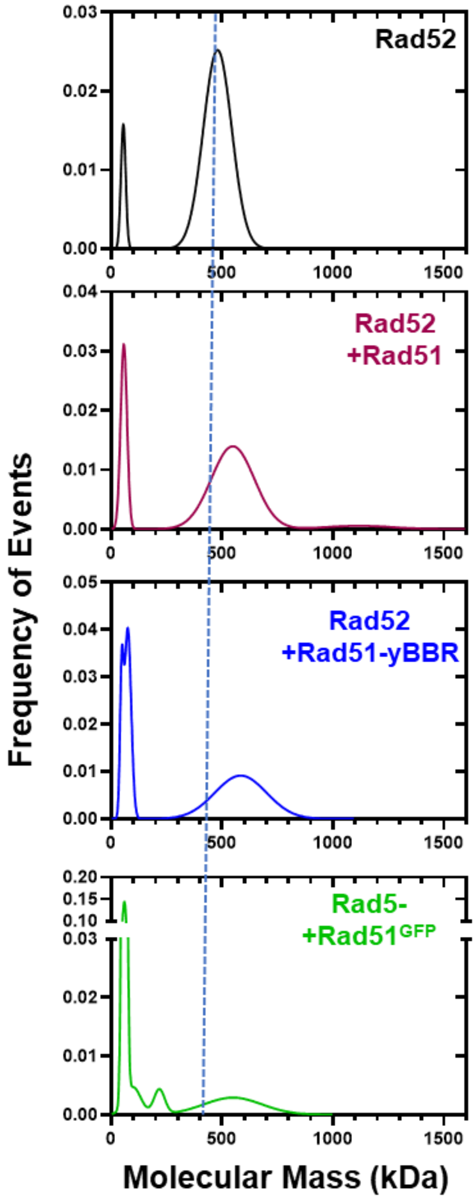


**Supplementary Figure 7. Fluorescent Rad51 protein interacts with Rad52.** Mass Photometry analysis of Rad52 interactions with various Rad51 variants are shown. Interactions between Rad52 and all forms of Rad51 used in this study are captured as noted by the shift in the Rad52 homodecameric peak position. The relative position of Rad52 alone is denoted by the dotted line. All experiments were done under conditions where all the Rad51 binding sites in Rad52 could be occupied (15 nM Rad52 homodecamer and 150 nM Rad51).

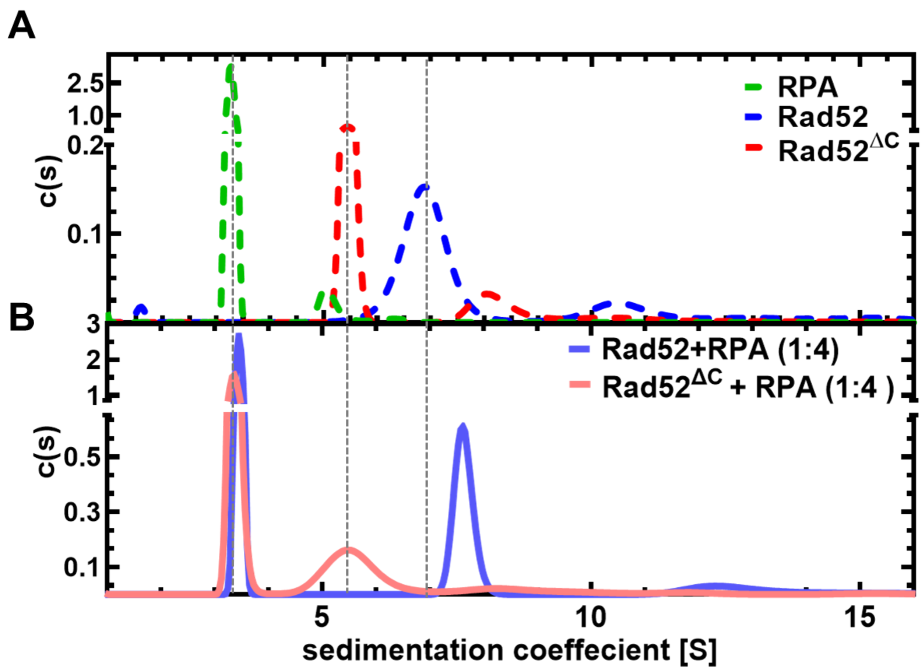


**Supplementary Figure 8. Analytical ultracentrifugation sedimentation velocity analysis of Rad52-Rad51 interactions.** Top panel shows the sedimentation profiles of RPA (green), Rad52 (blue), and Rad52$^{\Delta C}$ (blue). Bottom panel shows the sedimentation profiles of the RPA-Rad52 (blue) and RPA-Rad52$^{\Delta C}$ (pale pink) complexes. RPA interacts with Rad52, but does not bind to Rad52$^{\Delta C}$.

>sfGFP-Rad51

MSQVQEQHISESQLQYGNGSLMSTVPADLSQSVVDGNGNGSSEDIEATNGSGDGGGAGSAGGAGNRKRNG
MVSKGEELFTGVVPILVELDGDVNGHKFSVSGEGEGDATYGKLTLKFICTTGKLPVPWPTLVTTLTYGVQ
CFSRYPDHMKQHDFFKSAMPEGYVQERTIFFKDDGNYKTRAEVKFEGDTLVNRIELKGIDFKEDGNILGH
KLEYNYNSHNVYIMADKQKNGIKVNFKIRHNIEDGSVQLADHYQQNTPIGDGPVLLPDNHYLSTQSALSK
DPNEKRDHMVLLEFVTAAGITLGMDELYKGNRKRNGAGSAGGAGGGGLQEQAEAQGEMEDEAYDEAALGS
FVPIEKLQVNGITMADVKKLRESGLHTAEAVAYAPRKDLLEIKGISEAKADKLLNEAARLVPMGFVTAAD
FHMRRSELICLTTGSKNLDTLLGGGVETGSITELFGEFRTGKSQLCHTLAVTCQIPLDIGGGEGKCLYID
TEGTFRPVRLVSIAQRFGLDPDDALNNVAYARAYNADHQLRLLDAAAQMMSESRFSLIVVDSVMALYRTD
FSGRGELSARQMHLAKFMRALQRLADQFGVAVVVTNQVVAQVDGGMAFNPDPKKPIGGNIMAHSSTTRLG
FKKGKGCQRLCKVVDSPCLPEAECVFAIYEDGVGDPREEDE

Green: sfGFP sequence

Blue: additional flanking 16 aa linkers.

>ybbR-Rad51

MSQVQEQHISESQLQYGNGSLMSTVPADLSQSVVDGNGNGSSEDIEATNGSGDGDSLEFIASKLAGGLQE
QAEAQGEMEDEAYDEAALGSFVPIEKLQVNGITMADVKKLRESGLHTAEAVAYAPRKDLLEIKGISEAKA
DKLLNEAARLVPMGFVTAADFHMRRSELICLTTGSKNLDTLLGGGVETGSITELFGEFRTGKSQLCHTLA
VTCQIPLDIGGGEGKCLYIDTEGTFRPVRLVSIAQRFGLDPDDALNNVAYARAYNADHQLRLLDAAAQMM
SESRFSLIVVDSVMALYRTDFSGRGELSARQMHLAKFMRALQRLADQFGVAVVVTNQVVAQVDGGMAFNP
DPKKPIGGNIMAHSSTTRLGFKKGKGCQRLCKVVDSPCLPEAECVFAIYEDGVGDPREEDE

Pink: ybbR sequence

**Supplementary Figure 9. Sequence of the Rad51 variants used in this study.** The annotated amino acid sequences of sfGFP-Rad51 and ybbR-Rad51 constructs are shown.
